# PRDM16-DT: A Brain and Astrocyte-Specific lncRNA Implicated in Alzheimer’s Disease

**DOI:** 10.1101/2024.06.27.600964

**Authors:** Sophie Schröder, Ulrike Fuchs, Verena Gisa, Tonatiuh Pena, Dennis M Krüger, Nina Hempel, Susanne Burkhardt, Gabriela Salinas, Anna-Lena Schütz, Ivana Delalle, Farahnaz Sananbenesi, Andre Fischer

## Abstract

Astrocytes provide crucial support for neurons, contributing to synaptogenesis, synaptic maintenance, and neurotransmitter recycling. Under pathological conditions, deregulation of astrocytes contributes to neurodegenerative diseases such as Alzheimer’s disease (AD), highlighting the growing interest in targeting astrocyte function to address early phases of AD pathogenesis. While most research in this field has focused on protein-coding genes, non-coding RNAs, particularly long non-coding RNAs (lncRNAs), have emerged as significant regulatory molecules. In this study, we identified the lncRNA *PRDM16-DT* as highly enriched in the human brain, where it is almost exclusively expressed in astrocytes. *PRDM16-DT* and its murine homolog, *Prdm16os*, are downregulated in the brains of AD patients and in AD models. In line with this, knockdown of *PRDM16-DT* and *Prdm16os* revealed its critical role in maintaining astrocyte homeostasis and supporting neuronal function by regulating genes essential for glutamate uptake, lactate release, and neuronal spine density through interactions with the RE1-Silencing Transcription factor (Rest) and Polycomb Repressive Complex 2 (PRC2). Notably, CRISPR-mediated overexpression of *Prdm16os* mitigated functional deficits in astrocytes induced by stimuli linked to AD pathogenesis. These findings underscore the importance of *PRDM16-DT* in astrocyte function and its potential as a novel therapeutic target for neurodegenerative disorders characterized by astrocyte dysfunction

## Introduction

Astrocytes are abundant cell types in the central nervous system (CNS). Under physiological conditions, they provide trophic and metabolic support for neurons, and are crucial for synaptogenesis, synapse maintenance, and synaptic pruning. Additionally, they play roles in neurotransmitter recycling and maintaining the blood-brain barrier, ion, pH, and fluid homeostasis [1] [2] [3]. In response to CNS damage, such as injury or disease, astrocytes can undergo morphological, molecular, and functional changes, a process often referred to as ’reactive astrocytosis’ [4]. Depending on various factors, including the pathological context, affected astrocyte subpopulations, and environmental conditions, the adopted states can be either detrimental, where astrocytes contribute to neuroinflammatory processes while neuronal support function is compromised, or beneficial, acting in a neuroprotective manner [4].

There is increasing awareness of the role of astrocytes in neurodegenerative diseases such as Alzheimer’s disease (AD), as genes expressed in astrocytes have been genetically linked to AD pathogenesis. Moreover, atrophic and reactive astrocytes have been described in postmortem brain tissue from AD patients and in AD models [5] [6] [7] [8] [9]. Recent studies suggest a scenario in which activated microglia release interleukin 1a (IL1a), tumor necrosis factor alpha (Tnfa), and complement component 1q (C1q), driving astrocytes to a neurotoxic phenotype that contributes to synaptic dysfunction and neuronal cell death [3]. Similar results have been observed when astrocytes were exposed to amyloid beta 42 (Abeta24) [10] [11]. However, the molecular mechanisms underlying the formation of neurotoxic astrocytic states are only beginning to emerge. A better understanding of these processes could facilitate the development of therapeutic approaches towards neurodegenerative diseases.

In this context, basic and translational research in the past decades has mainly focused on the coding part of the genome, hence genes that are translated into proteins. However, only 1,5% of the human genome encodes for proteins while most of the transcriptome represents non-coding RNAs (ncRNAs), of which the majority is classified as long-non-coding RNAs (lncRNAs) [12].

lncRNAs are a heterogenous group of RNA molecules with a length of more than 300 nucleotides that lack the potential to code for proteins [13] [14]. Initially, many of these non-coding RNAs have been regarded as transcriptional noise. However, in the last years, it has become apparent that many lncRNAs are indeed functional RNA molecules that can play important roles in different biological processes [14] [15]. A significant proportion of lncRNAs exhibit tissue- and cell type-specific expression patterns, with approximately 40% of human lncRNAs reported to be brain-specific, implying their critical involvement in brain development and function [16], but most likely also in the establishment and progression of brain-related disorders [17]. These characteristics make them promising drug targets [18], especially considering the growing recognition of RNA therapeutics as a promising avenue for treating various diseases, including brain disorders [19] [20]. However, the knowledge about lncRNAs in the brain is still limited and especially for astrocytes, there is only very little information about functional involvements in the context of reactive astrocytosis.

In this study, we aimed to identify long non-coding RNAs (lncRNAs) that are specifically enriched in astrocytes and deregulated under conditions of reactive astrocytosis and in neurodegenerative diseases. We hypothesized that such lncRNAs could serve as *bona fide* candidates for targeted drug interventions. By combining single nucleus RNA sequencing (snucRNAseq) of the human brain with additional transcriptome data from human tissues we identify *PRDM16-DT* as the most highly enriched lncRNA in astrocytes with a murine homolog, named *Prdm16os*. *PRDM16-DT* is downregulated in reactive astrocytes and in the brains of individuals with AD. Knockdown of *Prdm16os* followed by total RNA sequencing and functional assays revealed its important role in maintaining astrocyte homeostasis and supporting neuronal function. Furthermore, we find that *Prdm16os* and *PRDM16-DT* regulate the expression of genes critical for synaptic support by functioning as a decoy for RE1-Silencing Transcription factor (Rest) in conjunction with the H3K27 methyltransferase Polycomb Repressive Complex 2 (PRC2). In line with these observations, reduced *Prdm16os and PRDM16-DT* levels affect glutamate uptake and lactate release which correlates with impaired neuronal cell viability and spine density. Finally, CRISPR-mediated overexpression of *Prdm16os* can mitigate functional deficits induced in reactive astrocytes. Overall, these results report for the first time a role of *Prdm16os* and *PRDM16-DT* in the brain and highlight its specific role in astrocyte function. Thus, *PRDM16-DT* is a potential novel therapeutic target for neurological disorders characterized by astrocyte dysfunction.

## Results

### Single Nucleus RNA Sequencing identifies *PRDM16-DT* as an astrocyte-specific lncRNA in the human brain

Our objective was to identify astrocyte-specific lncRNAs enriched in the brain and dysregulated in neurodegenerative disorders **(Fig. 1A)** . We hypothesized that studying such lncRNAs functionally could provide insights into the role of astrocytes in these diseases and potentially uncover promising drug targets with minimal side effects, given their brain and cell type-specific expression. To achieve this goal, we generated snucRNAseq data from postmortem brain tissues of healthy individuals using and adapted an iCell8 snucRNAseq protocol (Takara) with random primers, enabling us to detect both polyA and non-polyA transcripts. We generated over 400 million reads per sample and identified 17031 lncRNAs in total **(Fig. 1B)** . Next, we identified astrocyte-enriched lncRNAs by comparing the mean expression in astrocytes with that in all other cell types, resulting in ten lncRNAs with a ratio of >20 fold enrichment **(Fig. 1C)**. To facilitate functional analysis we decided to initially focus on lncRNAs (marked in red) that have a mouse homolog, allowing for further studies in both murine and human cells. LncRNAs were considered homologs when the genes were syntenically located in the same genomic locus, and the transcripts exhibited a sequence similarity of at least 40%. The most enriched and conserved lncRNA in astrocytes was *PRDM16-DT* which was almost exclusively expressed in astrocytes **(Fig. 1C-D)** . Next, we reanalyzed an RNAseq dataset encompassing 45 human tissues [21] and found that *PRDM16-DT* was significantly enriched in the central nervous system compared to other human tissues **(Fig. 1E)**.

**Figure 1:**
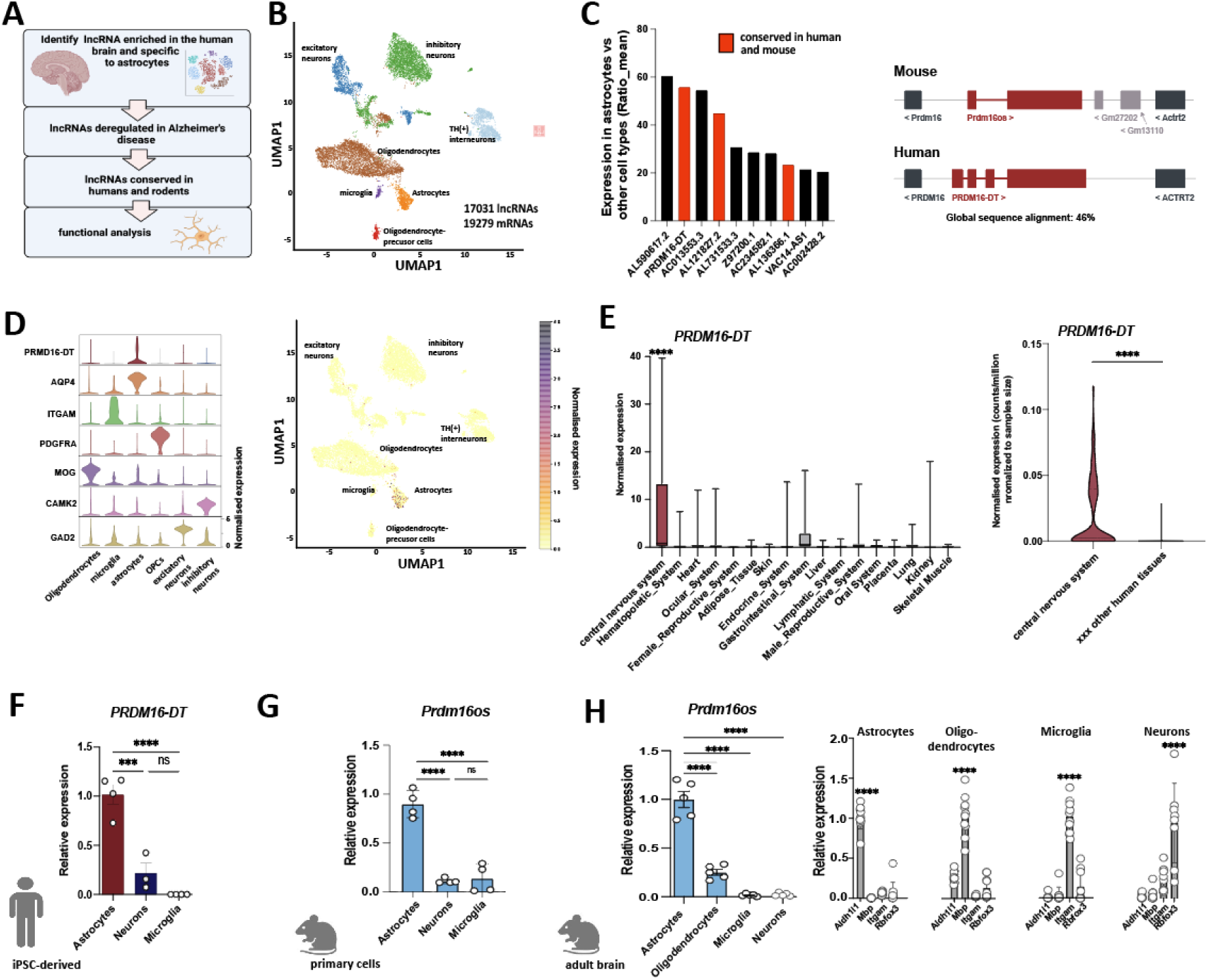
*PRDM16-DT/Prdm16os* is an astrocyte-specific lncRNA enriched in the brain. **A.** Strategy to identify astrocyte-specific lncRNAs in the brain and characterize their function. **B.** UMAP clustering based on snucRNAseq (iCELL8 technology) from healthy human brains (prefrontal cortex, BA9). **C.** Left panel: Bar graph showing the list of lncRNAs enriched in astrocytes when compared to all other cell types in the human brain. lncRNAs marked in red have a mouse homolog. Right panel: Schematic illustration showing the genomic localization of human *PRMD16*-DT and mouse *Prdm16os*. **D.** Left panel: Violin plot showing the expression of *PRDM16-DT* in different cell types of the human prefrontal cortex along with corresponding cell type marker genes. .Right panel: UMAP clustering as depicted in (B) showing the expression of *PRMD16-DT*. **E.** Left panel: Expression of PRDM16-DT in different human tissues (depicted are only tissues in which expression was detectable). One-way ANOVA revealed a significant difference among the groups (p < 0.0001). \*\*\*\**P* < 0.0001 for central nervous system vs. any of the other tissues (unpaired t-Tests). Right panel: Violin plot showing the expression of PRDM16-DT as counts per million normalized to the samples size for the central nervous system vs. the average expression across 44 other human tissues (only tissues in which PRDM16-DT expression was detected are shown) \*\*\*\**P* < 0.0001; unpaired t-Test. **F.** Bar chart showing the expression of *PRDM16-DT* in human iPSC-derived astrocytes, neurons and microglia (unpaired t test; ***P < 0.001, ****P < 0.0001, ns = not significant). **G.** *Prdm16os* expression in mouse primary astrocytes, neurons and microglia (unpaired t test; ***P < 0.001, ****P < 0.0001, ns = not significant). **H.** *Prdm16os* expression in astrocytes, oligodendrocytes, microglia and neuronal fraction isolated from the adult mouse brain (unpaired t test; ***P < 0.001). Right panel: Bar charts showing qPCR data from astrocytes, oligodendrocytes, microglia and neuronal fraction isolated from the adult mouse brain for marker genes for astrocytes (*Aldehyde Dehydrogenase 1 Family Member L1, Aldh1l* ), oligodendrocytes (*myelin basic protein, Mbp*), microglia (*Integrin Subunit Alpha M, Itgam* ) and neurons (*RNA Binding Fox-1 Homolog 3* , Rbfox3). Error bars indicate SD.

In summary, these data suggest that *PRDM16-DT* is a brain-enriched lncRNA predominantly expressed in astrocytes. To further validate this observation, we examined the expression of *PRDM16-DT* in human iPSC-derived cortical neurons, microglia, and astrocytes using qPCR. Consistent with the snucRNA-seq data, *PRDM16-DT* showed significant enrichment in astrocytes **(Fig. 1F)**. To assess whether this cell-type specificity is conserved across species, we analyzed the expression of the *PRDM16-DT* mouse homolog, *Prdm16os*, in primary astrocytes, neurons, and microglia **(Fig. 1G)** . Similar to human *PRDM16-DT*, mouse *Prdm16os* exhibited high enrichment in astrocytes. Furthermore, we employed MACS sorting to isolate astrocytes, oligodendrocytes, microglia, and a neuron-enriched fraction from the adult mouse brain. Consistent with our previous findings, *Prdm16os* was found to be most highly expressed in astrocytes **(Fig. 1H)**.

### *Prdm16os* and *PRDM16-DT* are downregulated in response to pathological insults and in Alzheimer’s disease

To investigate whether *PRDM16-DT* is dysregulated in neurodegenerative diseases, we analyzed its expression in postmortem brain samples (prefrontal cortex, BA9) obtained from healthy controls and patients with AD. Our analysis revealed a significant decrease in *PRDM16-DT* expression in AD patients **(Fig. 2A)** . This finding was further validated using data from the Agora database (https://agora.adknowledgeportal.org/), which includes over 1000 postmortem brain samples from both control individuals and AD patients. *PRDM16-DT* expression was found to be significantly reduced in the frontal pole and parahippocampal gyrus regions **(Fig. 2B)**.

**Figure 2:**
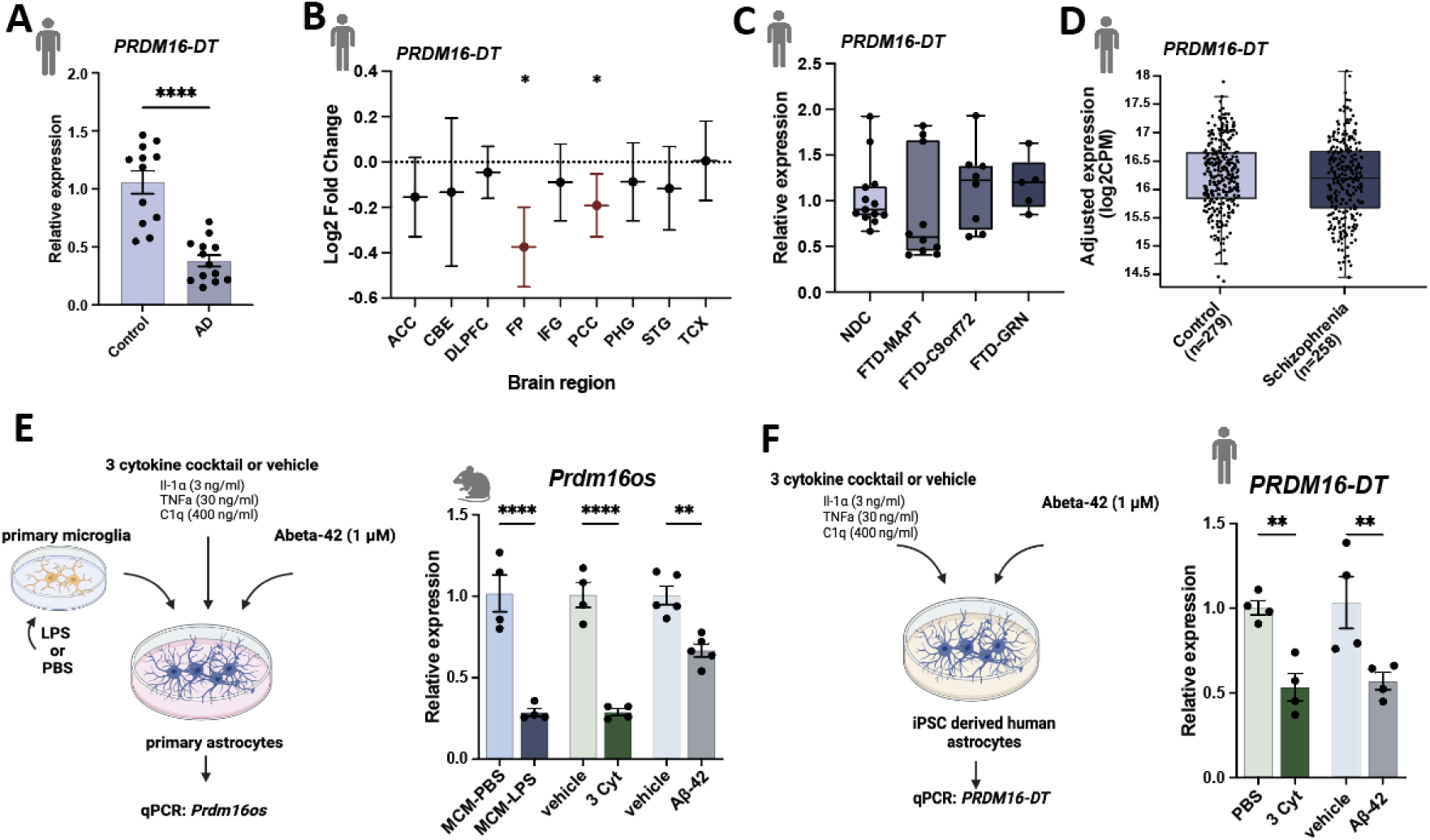
*Prdm16-DT* is decreased in the brains of AD patients and in response to AD risk factors. **A.** Bar chart showing qPCR data on the expression of *PRDM16-DT* in postmortem brain samples (prefrontal cortex, BA9) from control (n = 12) and AD patients (n = 13) (\*\*\*\**P* < 0.0001, unpaired t-Test) **B.** Log2 Fold changes of *PRDM16-DT* expression in different brain regions in AD patients compared to controls based on data from the Agora database (https://agora.adknowledgeportal.org/). (\**P* < 0.05). **C.** Bar chart showing the expression of PRDM16-DT in postmortem tissue samples (frontal lobe) of FTD patients with *MAPT* (n = 10), *C9ORF72* ( n = 8) or *GRN* (n = 6) mutations compared to non-demented controls (NDC, n = 13). **D.** Bar chart showing the expression of PRDM16-DT in postmortem brain tissue of controls (n = 279) compared to schizophrenia patients (n = 258) obtained from a study by Wu et al., [24]. **E.** Left panel: Experimental design. Right panel: Bar plot showing *Prdm16os* expression in mouse astrocytes after treatment with LPS-activated microglia conditioned medium (MCM-LPS) compared to control (MCM-PBS), a 3 cytokine cocktail (3 Cyt) and Aß-42 treatment compared to the corresponding vehicle controls.(\*\*\*\**P* < 0.00001, \*\**P* < 0.001 unpaired t-Test). **F.** Left panel: Experimental design. Right panel: Bar plot showing *PRMD16-DT* expression in human iPSC-derived astrocytes after treatment with a 3 cytokine cocktail (3 Cyt) and Aß-42 compared to the corresponding controls (\*\**P* < 0.01 unpaired t-Test). ACC: Anterior Cingulate Cortex, AD: Alzheimer’s Disease, CBE: Cerebellum, 3 Cyt: 3 cytokine cocktail, DLPFC: Dorsolateral Prefrontal Cortex, FTD: Frontotemporal Dementia, FP: Frontal pole; IFG: Inferior Frontal Gyrus, PCC: Posterior Cingulate Cortex, PHG: Parahippocampal Gyrus, STG: Superior Temporal Gyrus, TCX: Temporal Cortex. Error bars represent SD.

Subsequently, we analyzed *PRDM16-DT* expression in postmortem brain tissues (frontal lobe) obtained from patients with frontotemporal dementia (FTD) carrying mutations in *C9ORF72*, *GRN*, or *MAPT* genes, as well as from control individuals that are available via the RiMOD database [22] [23]. We did not observe any significant difference in *PRDM16-DT* expression between these groups **(Fig. 2C)**. Additionally, we examined *PRDM16-DT* expression in a transcriptome dataset derived from postmortem brains of individuals with schizophrenia, as compared to controls [24]. Similar to our findings in FTD patients, there was no discernible difference in *PRDM16-DT* expression between schizophrenia patients and controls **(Fig. 2D)** . These results collectively suggest that *PRDM16-DT* expression is decreased specifically in the brains of AD patients.

It is widely believed that the accumulation of amyloid beta (Aß) oligomers is among the earliest pathological changes in AD pathogenesis, occurring decades before clinical symptoms manifest [25] [26]. Reactive astrogliosis is also observed very early in mouse models for amyloid pathology [27]. While Abeta42 can directly affect astrocytes, a prominent mechanism by which Abeta42 affects astrocytes is via the activation of microglia [28], which in turn release pro-inflammatory cytokines IL-1α, TNF, and C1q that are sufficient to induce reactive astrogliosis [3]. Based on these findings, we wanted to test if the exposure of astrocytes to Aß oligomers, media from activated microglia, or inflammatory cytokines would affect the expression of *PRDM16-DT* and *Prdm16os*. Firstly, we assessed the expression of *Prdm16os* in primary astrocytes stimulated with microglia-conditioned medium (media from microglia treated with LPS; MCM-LPS), compared to control media (media from microglia treated with PBS; MCM-PBS) (Fig. 2E). We observed a significant decrease in Prdm16os levels in astrocytes following MCM-LPS treatment (Fig. 2F). Treatment of primary astrocytes with the of Il1α, TNF, and C1q (3 cytokine cocktail) also resulted in a significant decrease in Prdm16os levels, similar to the effect of microglia-conditioned media (Fig. 2E). Similarly, treating primary astrocytes with Abeta42 led to a significant reduction in Prdm16os levels (Fig. 2E). Furthermore, stimulation of human iPSC-derived astrocytes with the 3 cytokine cocktail or Abeta resulted in a similar downregulation of PRDM16-DT as observed in mouse astrocytes (Fig. 2F).

In summary, our data demonstrate that PRDM16-DT is reduced in the brains of AD patients. Additionally, exposure of both mouse and human astrocytes to well-established AD risk factors (MCM, 3 cytokine cocktail, Abeta42) results in a similar decrease in PRDM16-DT and Prdm16os levels. These findings provide a suitable experimental framework for studying the role of PRDM16-DT/Prdm16os in the context of AD.

### *Prdm16os* regulates genes involved in synaptic function, apoptosis and inflammatory processes

Given the lack of previous studies on *PRDM16-DT/Prdm16os* in the CNS, we aimed to investigate its role at the cellular and functional level. Since the function of lncRNAs is closely tied to their subcellular localization [29], we initially employed murine models to examine the localization of *Prdm16os*. Through a combination of RNAscope and immunofluorescence staining for *Prdm16os* and the astrocyte marker Gfap, we determined that *Prdm16os* is predominantly located in the nucleus of astrocytes in the adult mouse brain **(Fig. 3A)** . To validate this finding, we conducted nuclear and cytoplasmic fractionation of primary mouse astrocytes and compared the levels of *Prdm16os* in each compartment. Our analysis revealed that *Prdm16os* primarily resides in the nuclear fraction **(Fig. 3B)**, suggesting a potential role for *Prdm16os* in gene transcription regulation.

**Figure 3:**
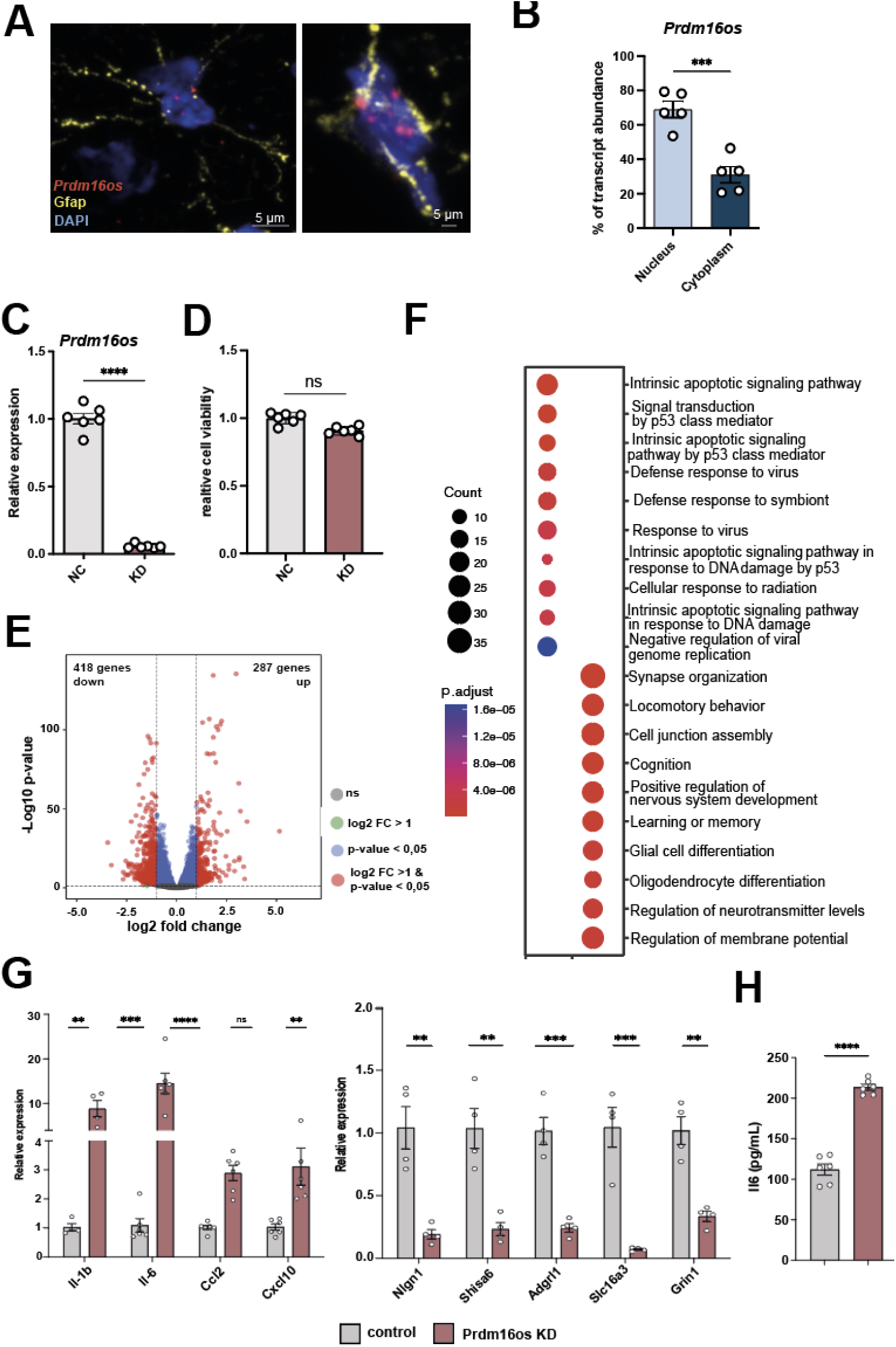
*Prdm16os* is localized to the nucleus and controls gene expression. **A.** Representative images from the adult mouse hippocampus showing the RNAscope signal for *Prdm16os* and immunofluorescence for Gfap. Nuclei are stained with DAPI. The right panel shows a higher magnification from a different hippocampal region. **B.** Bar plot showing qPCR values for *Prdm16os* in nuclear and cytoplasmic fractions prepared from primary astrocytes (\*\*\**P* < 0.001, unpaired tTest). **C.** Bar plot showing qPCR results for *Prdm16os* after treating primary astrocytes with GapmeRs to knock down (KD) *Prdm16os.* RNA was collected 48 hours after the addition of *Prdm16os GapmeRs* or control oligonucleotides (\*\*\*\**P* < 0.0001, unpaired tTest) **D.** Bar plot showing cell viability of primary astrocytes 48h after *Prdm16os* knock down in comparison to the treatment with control oligomers. **E.** Volcano plot shows the up-and down-regulated genes in primary astrocytes 48h after *Prdm16os* knock down. **F.** Plot showing the results of a GO term analysis for the up- and downregulated genes displayed in (E) (Analysis was done using clusterProfiler (v4.6.0) [30]. Two-sided hypergeometric test was used to calculate the importance of each term and the Benjamini-Hochberg procedure was applied for the P value correction). **G.** Bar plots showing the results of qPCR experiments for selected genes that were found to be deregulated upon *Prdm16os* knock down via RNAseq. Left panel: Selected up-regulated genes. Right panel: Selected downregulated genes \*\*\*\**P* < 0.0001, \*\*\**P* < 0.001; \*\**P* < 0.01; unpaired tTest). **H.** Bar chart showing the effect of Prdm16 knock down on IL-6 levels in the corresponding media measured via ELISA (\*\*\*\**P* < 0.0001, unpaired tTest). KD: knockdown, NC: negative control. Error bars indicate SEM.

With this insight, we proceeded to perform a knockdown (KD) of *Prdm16os* in primary astrocytes using a *Prdm16os* GapmeR probe **(Fig. 3C)**. We confirmed that *Prdm16os* knockdown had no adverse effects on cell viability after 48 hours **(Fig. 3D)** , prompting us to collect RNA samples for total RNA sequencing at this time point. Subsequent differential expression analysis unveiled significant changes, with 287 genes upregulated and 418 genes downregulated (log2FC > 1, FDR < 0.05, basemean > 50) **(Fig 3E, supplemental table 1)** . Gene Ontology (GO) term analysis of the up-regulated genes revealed pathways related to apoptosis such as “Intrinsic apoptotic signaling pathway” and inflammatory processes such as “Defense response to virus” **(Fig. 3F, supplemental table 2)** . The downregulated genes were linked to processes associated with synapse support function such as “synapse organization”, “cognition”, “learning and memory” and “regulation of neurotransmitter levels” **(Fig. 3F, supplemental table 2)** . We selected several of the upregulated genes linked to inflammation and downregulated genes linked to synaptic function to confirm the RNAseq data via qPCR **(Fig 3G)** . To test whether the increased expression of genes linked to inflammation would result in elevated secretion of the corresponding cytokines, we also measured IL-6 levels in the media of primary astrocytes treated with Prdm16os GapmerRs or control using ELISA. IL-6 levels were significantly increased (Fig 3H).

### The knockdown of *Prdm16os* in astrocytes leads to impaired glutamate uptake, lactate secretion and affects neuronal activity and spine density

The RNAseq data hinted at several cellular processes controlled by *Prdm16os*. For instance, the sequencing data suggested that the loss of *Prdm16os* may compromise the neuronal support function of astrocytes and we decided to study this potential mechanics further at first. One crucial role of astrocytes is to remove glutamate from the extracellular space. This process is primarily mediated by two main glutamate transporters, Glt-1 and Glast. Both glutamate transporters showed downregulation in the RNAseq dataset, a finding confirmed by qPCR analysis **(Fig. 4A)** and at the protein level **(Fig. 4B)**. Consistently, the uptake of extracellular glutamate was impaired following the knockdown (KD) of *Prdm16os* in primary astrocytes **(Fig. 4C)**.

**Figure 4:**
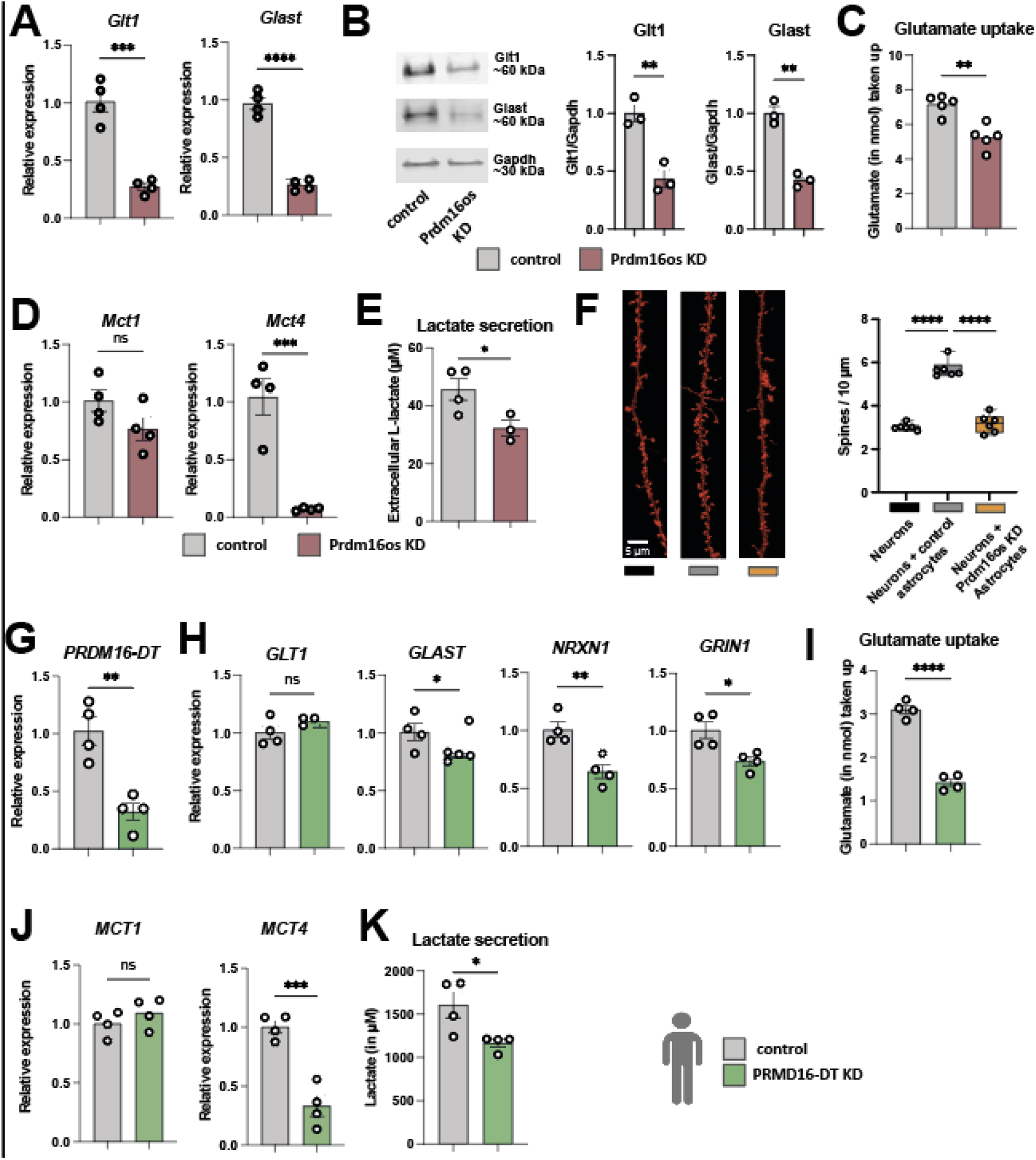
The loss of *Prdm16os* in astrocytes affects glutamate uptake, lactate secretion and neuronal function. **A.** qPCR showing the levels of the two glutamate transporters *Glt-1* and *Glast* after the KD of *Prdm16os* in primary astrocytes. **B.** Left panel: Representative Immunoblot images of Glt-1 and Glast after the KD of *Prdm16os* in astrocytes. Right panel: Quantification of (B) n = 3/group. **C.** Glutamate uptake after the KD of *Prdm16os* in astrocytes. **D.** qPCR showing the levels of the two lactate transporters *Mct1* and *Mct4* after the KD of *Prdm16os* in astrocytes. **E.** Lactate secretion after the KD of *Prdm16os* in astrocytes. **F.** Left panel: Representative images of dendrite and spine labeling of neurons cultured alone (Neurons) or with either control astrocytes (Neurons + control astrocytes) or *Prdm16os* KD astrocytes (Neurons + Prdm16os KD astrocytes). Right panel. Quantification of spine density. **G-K** shows data from human iPSC-derived astrocytes upon the KD of *PRDM16-DT*. **G.** KD of *PRDM16-DT* in human iPSC-derived astrocytes. **H.** qPCR showing the levels of the two glutamate transporters *GLT-1* and *GLAST* and two synapse plasticity genes *NRXN1* and *GRIN1* after the KD of *PRDM16-DT*. **I.** Glutamate uptake after the KD of *PRDM16-DT*. **J.** qPCR showing the levels of the two lactate transporters *MCT1* and *MCT4* after the KD of *PRDM16-DT*. **K.** Lactate secretion after the KD of *PRDM16-DT*.KD: knockdown, NC: negative control. Statistical significance was assessed by a one-way ANOVA with Tukey’s post hoc test for (F) or a Student’s unpaired t test for the other graphs; *P < 0.05, **P < 0.01, ***P < 0.001; ****P < 0.0001, ns = not significant.

In addition to glutamate uptake, astrocytes provide metabolic support in the CNS by secreting lactate, which is an important energy substrate for neurons. Also in this case, there are two relevant transporters expressed in astrocytes that are essential for memory function, Monocarboxylate transporter (Mct) 1 and Mct4 [31]. We confirmed via qPCR that the mRNA levels of *Mct4*, but not *Mct1*, were decreased in response to the KD of *Prdm16os* **(Fig. 4D)**. Next, we directly addressed the question if loss of *Prmd16os* in astrocytes would affect neuronal function. For this, we cultured neurons alone or in co-culture with astrocytes treated before with either *Prmd16os* GapmerRs (*Prdm16os* KD astrocytes) or control oligomers (control astrocytes). We observed that co-culturing primary neurons with control astrocytes was, as expected, able to enhance dendritic spine density, while no such effect was observed when neurons were co-cultured with *Prdm16os* KD astrocytes **(Fig. 4F)**. We also performed a knockdown of the human homologue of *Prmd16os* - *PRDM16-DT -* in human iPSC-derived astrocytes **(Fig. 4G)**. Whereas in mouse astrocytes the transcripts encoding both glutamate transporters were downregulated, in human astrocytes only *GLAST* was affected, while the levels of *GLT-1* remained unchanged **(Fig. 4H)** . We could also confirm the down-regulation of synaptic plasticity genes *NRXN1* and *GRIN1* **(Fig. 4H).** The observed functional consequences were similar across species, as also the human astrocytes showed reduced glutamate uptake after the KD of *PRDM16-DT* **(Fig. 4I)** . Similarly, also the expression of the lactate transporter *MCT4* and the secretion of L-lactate was decreased **(Fig. 4K-L)** . These data indicate that the general function of *Prdm16os* and *PRDM16-DT* as being important for neuronal support is conserved across species.

In addition to impaired neuronal support function, the RNAseq data suggested that loss of Prdm16os induces an inflammatory response (see Fig. 3G, H). However, while iPSC-derived astrocytes exhibited an increased expression of pro-inflammatory cytokines upon stimulation with the 3 cytokine cocktail, the induction of pro-inflammatory genes could not be confirmed when Prdm16-DT was knocked-down in human iPSC-derived astrocytes (Fig. S1).

Although the reason for this discrepancy is unclear at present, it may suggest that the control of inflammatory processes is not conserved across species and we therefore we decided to not follow up on the potential role of *Prdm16os* and *Prdm16-DT* in inflammation.

### *Prdm16os* interacts with Rest and Suz12 to modulate the expression of genes critical for synapse organization and function

So far our analysis suggests that *Prdm16os* and *PRDM16-DT* affect astrocytic processes linked to neuronal support via gene-expression control. One mechanism by which lncRNAs could control gene expression is to orchestrate the availability of transcription factors or chromatin regulators to bind specific DNA regions, such as promoter regions. To explore this further, we revisited the RNAseq data from astrocytes upon *Prdm16os* KD and focused on the downregulated genes since these were mainly associated with neuronal support function. We conducted an enrichment analysis to identify transcription factors potentially controlling the expression of these downregulated genes. This analyses revealed Suz12, a subunit of the Polycomb Repressive Complex 2 (PRC2) that mediates gene-repression via histone 3 lysine 27 tri-methylation (H3K27me3), followed by RE1-Silencing Transcription factor (Rest), also known as Neuron-Restrictive Silencer Factor (Nrsf) **(Fig. 5A)**. Previous studies have shown that lncRNAs can recruit PRC2 to specific chromatin locations [32] [33] [34]. Moreover, it has been described that PRC2 and Rest interact to silence gene-expression [35] [36] [37], with Rest being a key transcriptional regulator that represses neuronal genes in non-neuronal cells [38]. At the same time it is known that due to their neuronal support function and under specific conditions, astrocytes express genes typically associated with neurons such as ion channels and genes associated with synaptic function including neurotransmitter receptors, cell adhesion molecules, and other neuromodulatory genes, thus contributing to synaptic modulation [38] [39] [40] [41]. Furthermore, astrocytes are capable of transferring RNA, proteins, and even mitochondria to neurons, promoting synaptic plasticity [42] [43] [44] [45]. It is therefore tempting to speculate that *Prdm16os* may affect the function of PRC2 and Rest to fine tune the expression of genes linked to neuronal support function in astrocytes.

**Figure 5:**
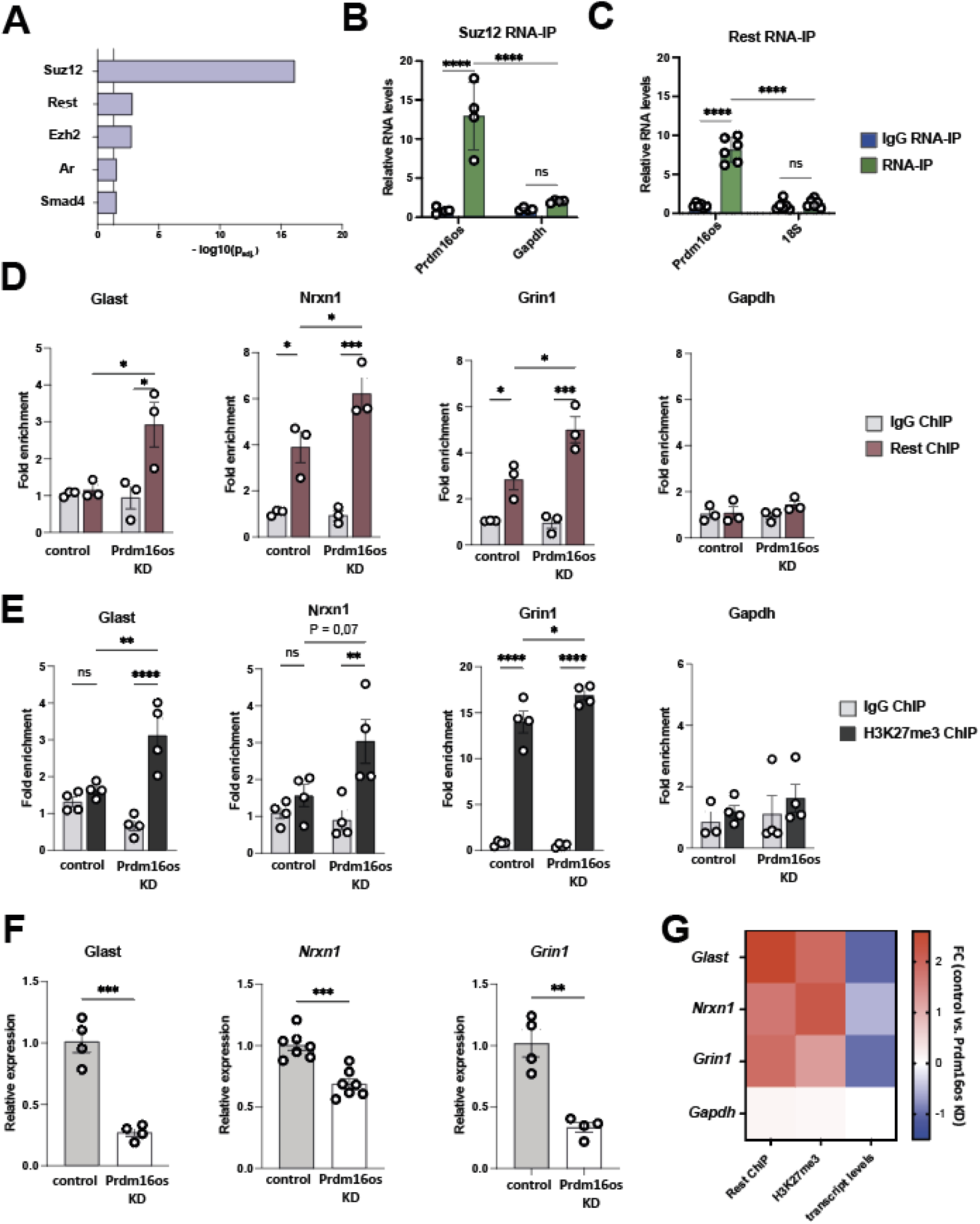
*Prdm16os* binds to Suz12 and Rest and affects Rest and H3K27me3 levels at the promoter of genes important for neuronal function b. **A.** Enrichment analysis for transcription factors based on the downregulated genes after the KD of *Prdm16os*. **B.** *Prdm16os* interacts with Suz12 as shown by RNA immunoprecipitation (RNA-IP). Gapdh mRNA was used as additional control. **C.** *Prdm16os* interacts with Rest as shown by RNA immunoprecipitation (RNA-IP) 18s RNA was used as additional control. **D.** ChIP for Rest of primary astrocytes treated with control oligomers (control) or *Prdm16os* GapmeRs (*Prdm16os* KD), followed by qPCR for promoter regions of genes important for neuronal support and synapse function (*Glast, Nrxn1, Grin1* ), which were downregulated upon *Prdm16os* KD. The housekeeping gene Gapdh is used as a control. **E.** ChIP for H3K27me3 of primary astrocytes treated with control oligomers (control) or *Prdm16os* GapmeRs (*Prdm16os* KD) followed by qPCR for promoter regions of the genes *Glast*, *Nrxn1* and *Grin1*. The housekeeping gene *Gapdh* is used as control. **F.** Bar charts showing the expression of *Glast*, *Nrxn1* and *Grin1* in astrocytes upon *Prdm16os* KD. **G.** Heat map showing the fold changes of Rest and H3K27me3 levels as well as the transcript levels of *Glast, Nrxn1, Grin1* and *Gapdh* in astrocytes upon *Prdm16os* KD. Statistical significance was assessed by a one-way ANOVA with Tukey’s post hoc test; *P < 0.05, ***P < 0.001; ****P < 0.0001, ns = not significant.

To test this hypothesis, we conducted RNA-immunoprecipitation (RNA-IP) for Suz12 **(Fig. 5B)** and Rest **(Fig. 5C)** and demonstrated their interaction with *Prdm16os*. To investigate whether *Prdm16os* affects the interaction Rest with promoter regions of potential target genes, we performed Suz12 chromatin immunoprecipitation followed by qPCR (ChIP-qPCR) in primary astrocytes that were treated with a control oligonucleotides (control) or GapmeRs to mediate the knockdown of *Prdm16os*. For qPCR, we selected genes for which we had confirmed their decreased expression upon *Prdm16os* and Prdm16-DT knockdown in primary astrocytes and iPSC-derived human astrocytes. We found that the levels of Rest binding to the promoter regions of candidate mRNAs linked to synaptic plasticity, namely *Glast*, *Nrxn1* and *Grin1* were increased after the KD of *Prdm16os*. There were no significant changes in Rest binding to the promoter of glycerinaldehyd-3-phosphat-dehydrogenase (*Gapdh*) that was used as a housekeeping control gene **(Fig. 5D)** .Since Suz12 is part of the PRC2 complex regulating H3K27me3, we decided to analyze if the KD of *Prdm16os* would affect H3K27me3, a mark linked to gene repression. ChIP-qPCR analysis from astrocytes treated with control oligomers or GapmeRs targeting *Prdm16os* revealed significantly increased H3K27me3 levels at the promoter regions of Glast and Grin1 upon *Prdm16os* KD **(Fig 5E)**. For *Nrx1*, we detected a non-significant trend, and no effect was observed for Gapdh **(Fig 5E)**. Finally, we could confirm for this experimental setting that the transcripts for *Glast, Nrxn1 and Grin1* were decreased in astrocytes upon *Prdm16os* knock down **(Fig 5D)**. These findings suggest that *Prdm16os* might act as a decoy for Rest and Suz12 to enable and fine-tune the expression of astrocytic genes that have important neuronal support functions.

### CRISPR-mediated restoration of *Prdm16os* expression can rescue cytokine-induced functional deficits in astrocytes

Given the downregulation of *Prdm16os* in the brains of AD patients and in astrocytes exposed to AD risk factors (see Fig. 2), our next objective was to investigate whether reinstating the expression of this lncRNA would be sufficient to ameliorate the observed functional defects. To achieve this, we decided to use the 3-cytokine cocktail exposure model (see Fig 2E) and employed CRISPR activation (CRISPRa) targeting the promoter region of *Prdm16os* via guide RNAs (gRNA). As a control, a gRNA with no target in the DNA was used (Scramble). We tested 3 different gRNAs under basal conditions and observed that gRNA-1, and 3 significantly increased the expression of *Prdm16os* with gRNA-3 leading to a 2-fold increase **(Fig. 6A).** Considering the superior efficacy of gRNA-3, it was selected for subsequent experiments. The plasmid to express CRISPRa also contained a Gfp signal allowing us to confirm successful transfection of primary astrocytes **(Fig. 6B)** . Next, astrocytes were transfected with CRISPRa and gRNA-3 (gRNA-3) upon concurrent stimulation with the 3 cytokine cocktail (3 Cyt), or vehicle as control (vehicle). Cell transfected with CRISPRa and scramble control guide RNA were used as control (Scramble) **(Fig. 6C_MODEL)**.

**Figure 6:**
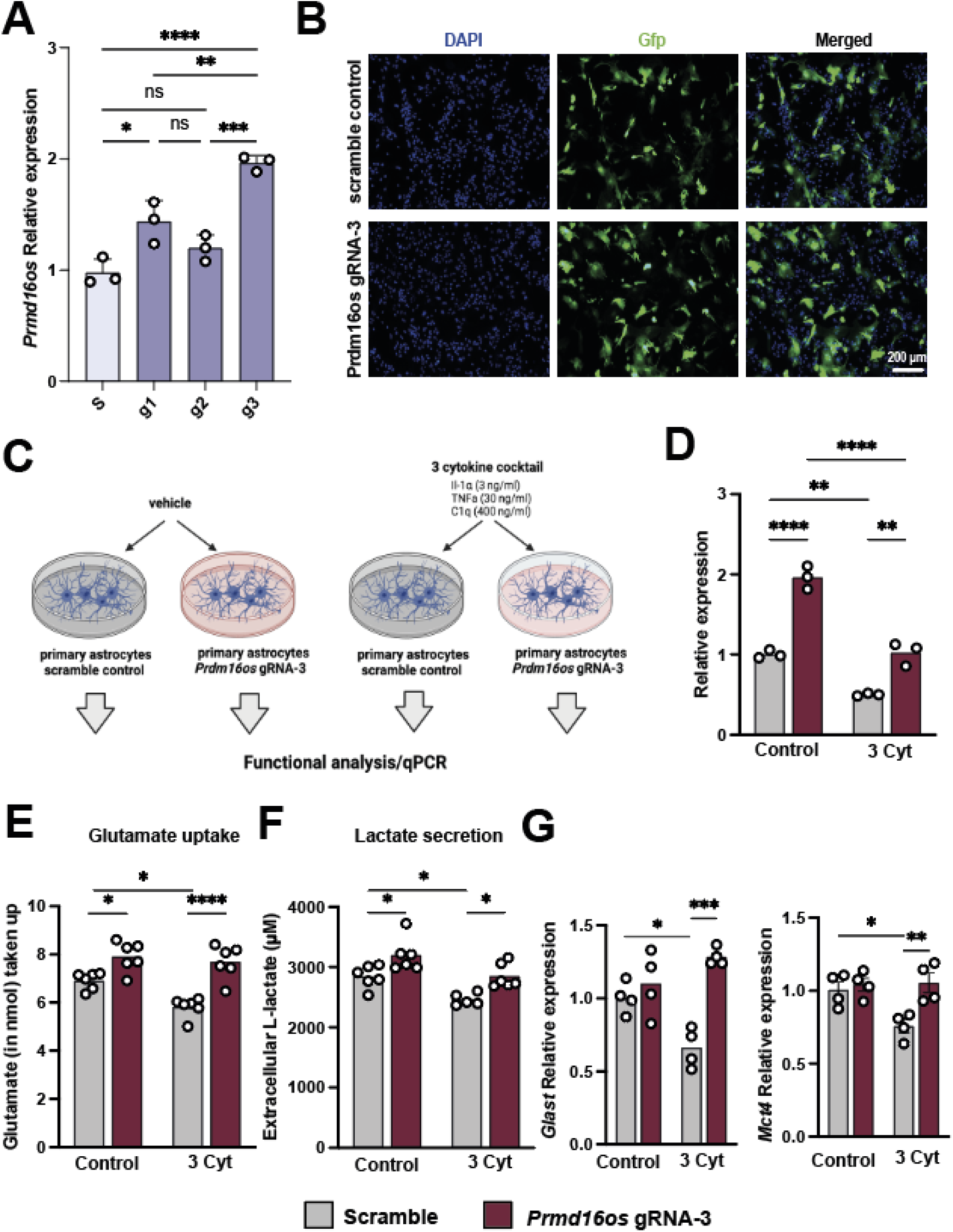
CRISPRa-mediated overexpression of *Prdm16os* can rescue functional impairments induced by cytokine treatment. **A.** Bar Chart showing the expression of *Prdm16os* in astrocytes transfected using CRRIPRa in combination with g1, 2 or 3 in comparison of cells treated with scrambled RNA (s). **B.** Representative immunofluorescence images showing the transfection of primary astrocytes with CRISPRa plasmids containing a scramble or *Prdm16os*-targeting g3 RNA. **C.** Scheme of the experimental approach. **D.** qPCR showing the levels of *Prdm16os* in astrocytes treated with either the 3 cytokine cocktail (3 Cyt) ot vehicle (control) in the presence of a either scramble RNA or g3 RNA to mediate CRISPRa of *Prdm16os.* Note that CRISPRa with g3 increased *Prdm16os* expression in the control group when compared to scramble RNA (one-way ANOVA followed by tTest; **P < 0.01) and is able to reinstate *Prdm16os* expression to physiological levels upon cytokine treatment (one-way ANOVA followed by tTest; ***P < 0.001). **E.** Bar charts showing the results from the Glutamate uptake assay of primary astrocytes after cytokine treatment and CRISPRa. **F.** Bar charts showing the results from the Lactate release assay of primary astrocytes after cytokine treatment and CRISPRa. Note that cytokine treatment impairs glutamate uptake and lactate release which is reinstated upon CIRPRa mediated overexpression of *Prdm16os.* **G.** Bar charts showing qPCR results for *Glast* (upper panel) and *Mct4* (lower panel) in the 4 experimental groups. Note that CRIPRa mediated overexpression can reinstate physiological expression of *Glast* and *Mct4* upon cytokine treatment. For A, D-G: Statistical significance was assessed by a one-way ANOVA with Tukey’s post hoc test; *P < 0.05, **P < 0.01; ****P < 0.0001

In line with our previous observations, treating cells with the 3 cytokine cocktail reduced the levels of *Prdm16os*, while CRISPRa treatment reinstated *Prdm16os* expression to physiological levels **(Fig. 6D)**. Encouraged by this observation, we wanted to test if the reinstatement of *Prdm16os* expression would also affect glutamate uptake and lactate release, which were both reduced upon *Prdm16os* KD and are compromised in AD pathogenesis. In agreement with this, glutamate uptake **(Fig. 6E)** and lactate release **(Fig. 6F)** was significantly impaired upon cytokine treatment. However, when astrocytes were treated with cytokines when *Prdm16os* expression levels were increased via CRISPRa, this was sufficient to reinstate glutamate uptake **(Fig. 6C)** and lactate release **(Fig 6D)** .

Interestingly, glutamate uptake and lactate release was also increased when *Prdm16os* levels had been elevated via CRIPRa in vehicle-treated cells **(Fig. 6E,F)** further suggesting a key role of *Prdm16os* in the regulation of these vital astrocyte functions. Using qPCR we confirmed that key genes linked to glutamate uptake (*Glast*) and lactate release (*Mct4*) that we had previously found to be affected by *Prdm16os* and *PRDM16-DT* KD were significantly decreased upon 3 Cyt treatment (**Fig 6G)**. CRISPR a-mediated increased expression of *Prdm16os* was able to ameliorate this effect **(Fig 6G)**.

In conclusion these data suggest that strategies to elevate *Prmd16-DT* could help to reinstate important neuronal support functions of astrocytes that are compromised in response to stimuli associated with AD pathogenesis.

## Discussion

In this study, we identified the human lncRNA *PRDM16-DT* and its mouse homologue, *Prdm16os*, to be enriched in the brain, where they are almost exclusively expressed in astrocytes when we cortical tissues. This finding aligns with previous studies demonstrating that lncRNAs can exhibit tissue-specific expression patterns [16]. We focused on *PRDM16-DT* in this study because it also has a homolog in mice, and conservation between species is an indicator of functionality [46]. Further analysis revealed that *PRDM16-DT* levels are decreased in the brains of Alzheimer’s disease (AD) patients, while no change in expression was observed in schizophrenia patients or in the brains of patients with frontotemporal dementia (FTD). Although data on lncRNAs in the brains of AD patients are still rare compared to the analysis of the coding transcriptome [47] [48], previous studies have reported lncRNAs deregulated in postmortem brains from AD patients. For example, *BACE1-AS*, which controls the production of amyloid beta peptides, has been reported to be deregulated [49].

While analyzing human brain tissue provides a unique opportunity to elucidate the processes underlying AD pathogenesis, the data might be affected by factors such as post-mortem delay and other variables [50] [51]. Therefore, it is important to note that *PRDM16-DT* and *Prdm16os* expressions were also decreased when human iPSC-derived or mouse astrocytes were exposed to stimuli associated with AD. These stimuli included microglia-conditioned media after LPS treatment, a 3-cytokine cocktail shown to mediate microglia-induced reactive astrogliosis [3], and Abeta42.

In line with the observation that *Prdm16os* was mainly localized to the nucleus, we observed a substantial effect on gene expression upon knockdown of *Prdm16os*, with the expression of genes important for neuronal support being downregulated. Among these were the glutamate transporter *Glast* and the lactate transporter *Mct4*. Similarly, *GLAST* and *MCT4* were downregulated when *PRDM16-DT* was knocked down in human iPSC-derived astrocytes. Consistently, KD of *Prdm16os* or *PRDM16-DT* in mouse or human astrocytes impaired glutamate uptake and lactate release. Loss of *Prdm16os* expression in astrocytes also increased the expression of proinflammatory cytokines. However, this effect was not observed in human iPSC-derived astrocytes, despite their confirmed response to the 3-cytokine cocktail stimulation, as evidenced by increased expression of cytokines such as *IL6* and *IL1b*.

While the molecular mechanisms underlying this discrepancy remain unclear, it is interesting to note that studies have reported that during AD pathogenesis, astrocytes exhibit decreased levels of the glutamate transporters *Glast* and genes involved in synapse organization, such as neuroligins and neurexins [52] [53]. These genes were all downregulated upon *Prdm16os* or *PRDM16-DT* knockdown in our study. Moreover, snucRNAseq studies on the brains of AD patients suggest a modest increase in proinflammatory genes in astrocytes and, more strikingly, a strong decrease in the expression of homeostatic genes involved in synapse regulation [54]. This is in line with other data on familial AD cases suggesting that astrocytes in AD brains are characterized by transcriptional underexpression of key genes linked to astrocytic function [54].

Nevertheless, we cannot exclude a role for *PRDM16-DT* in inflammatory processes in human astrocytes during AD pathogenesis. Future research is necessary to elucidate this aspect in more detail.

Taken together, the data from our analysis of postmortem human brains and human and mouse astrocytes consistently suggest a role for astrocytic *Prdm16os* and *PRDM16-DT* in neuronal support functions. In line with this, we observed that astrocytes failed to enhance spine density in the absence of *Prdm16os*. Although there is no data for *PRDM16-DT* or *Prdm16os*, and limited data on the role of lncRNAs in astrocytes, our findings align with previous studies showing that lncRNAs can regulate astrocyte function. For example, *Neat1*, a ubiquitously expressed lncRNA, is increased in astrocytes in response to ischemia and AD models [55] [56]. Similarly, *MEG3* is a lncRNA upregulated in human neurons xenografted in AD mouse models, and its inhibition ameliorated amyloid-induced necroptosis [57]. This aligns with previous findings showing that *MEG3* inhibition could rescue Abeta-induced phenotypes in a rat model of AD, including reactive astrogliosis [58]. Other lncRNAs associated with neurodegenerative conditions and astrocyte function include *NKILA* and *MALAT1* [59] [60].

However, unlike *PRDM16-DT,* all these lncRNAs are ubiquitously expressed, and therefore targeting them therapeutically may lead to unwanted side effects. Therefore, targeting lncRNAs exclusive to the brain and specifically to astrocytes offers the potential for more precise therapeutics. Along this line, our data show that CRISPRa-mediated increase of *Prdm16os* expression in astrocytes can ameliorate the phenotypes induced by exposure to AD risk factors. These findings are significant because targeting astrocyte function has been suggested as a suitable approach for addressing specific early phases of AD pathogenesis [7].

Our data suggest that *PRDM16-DT* and *Prdm16os* may mainly influence astrocyte function by controlling gene expression. This aligns with previous studies indicating that a key role of lncRNAs is the regulation of gene expression in the nucleus, often through interactions with transcription factors or chromatin-modifying enzymes [61] [14]. We observed that *Prdm16os* interacts with Suz12, a subunit of the PRC2 complex that orchestrates gene silencing via H3K27-trimethylation [35] [36] [37], and Rest, a key transcriptional regulator that represses neuronal genes in non-neuronal cells [38]. This is interesting since Rest and PRC2 were shown to interact to control gene expression [35].

While Rest orchestrates the repression of neuronal genes in non-neuronal cells, astrocytes are known to express several such genes, including synaptic cell adhesion molecules like neurexins and neuroligins that mediate astrocyte-synapse interactions [62] and other synapse-associated genes [63] [64] [65]. It is tempting to speculate that a mechanism exists to prevent Rest from binding to the promoter regions of these genes in astrocytes, thereby preventing complete gene silencing. Our data suggest that *Prdm16os* may regulate the expression of these target genes by acting as a decoy for Rest and Suz12. In this scenario, the loss of *PRDM16-DT*, as observed in AD brains, would compromise the fine-tuning of gene expression pathways in astrocytes, leading to a loss of neuronal support function and synaptic plasticity. This view is consistent with previous data suggesting that impaired astrocyte function plays an important role in neurodegenerative processes [66] [28] [67] [68].

It is important to acknowledge the limitations of our study. Increasing evidence suggests astrocyte heterogeneity across the brain, revealing several distinct astrocyte subtypes between and within brain regions [69] [70]. Investigating the expression pattern of *Prdm16os* and *PRDM16-DT* in brain regions other than the cortex might provide further insights into their role in supporting neuronal function and in the course of AD. Another important question for future research is to elucidate the regulation of *PRDM16-DT* expression to better understand the mechanisms underlying its reduced levels in the brains of AD patients and in AD model systems. Although our data suggest that *PRDM16-DT* is specifically decreased in the brains of AD patients while remaining unaffected in FTD and schizophrenia patients, we cannot exclude the possibility that *PRDM16-DT* is affected in other brain diseases. Additionally, it could be considered to test the role of *Prdm16os* in transgenic mouse models. However, in line with the principles of the 3Rs (Replacement, Reduction, and Refinement), in this study we decided to focus on cellular models and confirm key findings in human iPSC-derived cells rather than mouse models with the aim to reduce the reliance on animal experimentation, which is getting increasingly difficult in the European Union and in Germany in particular.

In summary, our data suggest that *PRDM16-DT* could be a suitable drug target to ameliorate neurodegenerative processes associated with AD pathogenesis. The specific expression pattern of *PRDM16-DT* in astrocytes of the human brain offers a unique possibility to target distinct phases of AD pathogenesis with minimal unwanted side effects.

## Methods and Material

### Human tissues

For snucRNAseq tissue samples (prefrontal cortex, BA9) were obtained with ethical approval from the ethics committee of the University Medical Center Göttingen and upon informed consent from the New York brain bank (female: n = 1, age = 54, postmortem delay: 6:41h; male: n = 1; age = 62, postmortem delay = 5:24h). Brains (prefrontal cortex, BA9) from control (n = 4 females & 5 males; age = 73,7 ± 9,5 years, PMD = 20,5 ± 4,1 h) and AD patients (n = 4 females & 6 males; age = 85,3 ± 7,3 years, PMD = 16,4 ± 7,4 h; Braak & Braak stage IV) were obtained with ethical approval from the ethics committee of the University Medical Center Göttingen and upon informed consent from the Harvard Brain Tissue Resource Center (Boston, USA).

### Sorting of neuronal and non-neuronal nuclei from human brain

The isolation protocol of single nuclei from frozen human brain was adapted from the previously published protocol with following modifications [71] [72]. In short, frozen tissues were homogenized using a Dounce homogenizer in 500 µl EZ prep lysis buffer (Sigma NUC101) supplemented with 1:200 RNAse inhibitor (Promega, N2615) for 60 times in a 1.5 ml Eppendorf tube using micro pestles. The volume was adjusted to 2000 µl with lysis buffer and incubated on ice for 7 min. Cell lysates were centrifuged for 5 min at 500 × g at 4 °C and supernatants were discarded. The pellet was resuspended in 2000 µl lysis buffer followed by an incubation on ice for 5 min. Lysates were centrifuged (500 × g at 4 °C) and the pellet was resuspended in 1500 µl nuclei suspension buffer (NSB, 0.5% BSA, 1:100 Roche Protease inhibitor, 1:200 RNAse inhibitor in 1 × PBS) centrifuged again for 5 min (500 × g at 4 °C). The pellet was finally resuspended in 1000 µl NSB and stained with anti-NeuN-AlexaFluor®488 (MAB377X) for 1h at 4°C rotating followed by centrifugation for 5 min (500 × g at 4 °C). The pellet was washed once with 500 µL NSB and the pellet was resuspended in 300µL NSB and stained with 1:100 7AAD (Invitrogen, Cat: 00–6993-50). Single NeuN-positive and NeuN-negative nuclei were sorted using the BD FACS Aria III sorter. Sorted nuclei were counted in the Countess II FL Automated Counter. Single-nuclei RNA-seq using the iCELL8 System was performed at the NGS Integrative Genomics Core Unit in Göttingen, Germany. Approximately 1200 single nuclei per sample were sequenced.

### Single nucleus total RNA sequencing and Analysis

The Takara ICELL8 5,184 nano-well chip was used with the full-length SMART-Seq ICELL8 Reagent Kit for single nuclei library performances as described in [73]. Briefly, nuclei suspensions were fluorescent-labelled with live/dead stain, Hoechst 33,342 for 15 min prior to their dispensing into the Takara ICELL8 5,184 nano-well chip. CellSelect Software (Takara Bio) was used to visualize and select wells containing single nuclei. A total of four 5184 nano-wells chips were used for all samples obtaining 1200 to 1400 nuclei/sample. Final libraries were amplified and pooled as they were extracted from each of the single nanowell chip. Pooled libraries were purified and size selected using Agencourt AMPure XP magnetic beads (Beckman Coulter) to obtain an average library size of 500 bp. A typical yield for a library comprised of ∼ 1,300 cells was ∼ 15 nM. Libraries were sequenced on the HiSeq 4000 (Illumina) to obtain on average of 0.3 to 0.4 Mio reads per nuclei (SE; 50 bp). Illumina’s conversion software bcl2fastq (v2.20.2) was used for adapter trimming and converting the base calls in the per-cycle BCL files to the per-read FASTQ format from raw images. For further processing, the Cogent NGS Analysis pipeline (v1.5.1) was used to generate a gene-count matrix. The demuxer (cogent demux) was used to create demultiplexed FASTQ files from the barcoded sequencing data. The resulting output was then used as input for the analyzer (cogent analyze) which performs trimming, mapping and gene counting to create a gene counts matrix. Quality control was done by evaluating the quality report provided by the Cogent analyzer.

The SCANPY package was used for pre-filtering, normalization and clustering. Initially, cells that reflected low-quality cells (either too many or too few reads, cells isolated almost exclusively, cells expressing less than 10% of house-keeping genes) were excluded. Next, counts were scaled by the total library size multiplied by 10.000, and transformed to log space. Highly variable genes were identified based on dispersion and mean, the technical influence of the total number of counts was regressed out, and the values were rescaled. Principal component analysis (PCA) was performed on the variable genes, and UMAP was run on the top 50 principal components (PCs). The top 50 PCs were used to build a k-nearest-neighbours cell–cell graph with k= 50 neighbours. Subsequently, spectral decomposition over the graph was performed with 50 components, and the Leiden graph-clustering algorithm was applied to identify cell clusters. We confirmed that the number of PCs captures almost all the variance of the data. For each cluster, we assigned a cell-type label using manual evaluation of gene expression for sets of known marker genes by plotting marker gene expression on the UMAP and visual inspection. Violin plots for marker genes were created using the “stacked_violin function” as implemented in SCANPY.

### Primary astrocyte culture

Primary mouse astrocyte cultures were prepared from postnatal day 1-2 mice as previously described [74] with minor modifications. Briefly, pups were sacrificed by decapitation and the brains quickly removed. Cortices and hippocampi were dissected and dissociated using 0.05 % Trypsin-Ethylenediaminetetraacetic acid (EDTA) (Gibco). Cells from 2-3 mice were plated into one T75 flask coated with poly-D-lysine (PDL) and cultured with DMEM, 10% fetal bovine serum and 1% penicillin/streptomycin (all Gibco) (DMEM+) in a humidified incubator with 5% CO2 at 37°C for 7-8 days. Then, cells were placed on a shaker (Incu-Shaker Mini, Benchmark) and shaken at 160 rpm for 6 hours to loosen neurons and non-astrocytic glia. The medium was removed and the astrocytes incubated with 0.25% Trypsin-EDTA (Gibco) for 4 minutes at 37°C. DMEM+ was added to inactivate the trypsin and the cell suspension was transferred to fresh tubes and centrifuged for 4 minutes at 400 g. The cells were resuspended in Neurobasal™ Plus Medium containing 2% B-27™ Plus Supplement, 1% penicillin/streptomycin, 1x GlutaMAX™ Supplement (all from Gibco) (NB+) and 5 ng/mL Heparin-Binding EGF-Like Growth Factor (HB-EGF) (Sigma-Aldrich) and plated at a density of 15,000 cells/cm^2^ onto PDL- or 0.1% polyethylenimine (PEI)-coated cell culture dishes. Cultures were then maintained in the incubator, and half the medium was changed two times a week.

### Primary cortical neuron culture

Pregnant CD-1 mice were sacrificed using pentobarbital. Embryonic day 17 embryos were taken out and the brains quickly removed in ice-cold PBS. The meninges were removed and the cortices dissected. Then, cell dissociation was performed using the Papain Dissociation System (Worthington) as described by the manufacturer’s instructions. The cells were resuspended in NB+ and seeded at a density of 120,000 cells/cm^2^ for glutamate treatment or 60,000 cells/cm^2^ for spine density analysis.

For the co-cultures, inserts containing astrocytes were added to cortical neurons on day in vitro (DIV) 14 and the experiments were performed on DIV17.

### Human iPSC-derived astrocytes

Human iPSC-derived astrocytes (Ncyte Astrocytes) were obtained from Ncardia. Cells were thawed and cultured according to the manufacturer’s instructions. After thawing, the astrocytes were left in culture for 1-2 weeks before performing experiments.

### Stimulation of primary astrocytes

Primary microglia were cultured as previously described [75]. On DIV 10, they were stimulated with 100 ng/mL Lipopolysaccharide (LPS) for 4 hours. Then, the medium was changed to NB+, collected after 6 hours, filtered using a 0.22 µm filter and stored at -80°C until use. On DIV13 of astrocyte culture, it was added as a 1:1 mixture and RNA was isolated 24 hours later.

To mimic the activation of astrocytes by microglia, primary astrocytes were treated with a cytokine mixture consisting of Il-1α (3 ng ml−1, Sigma, I3901), TNF (30 ng ml−1, Cell Signaling Technology, 8902SF) and C1q (400 ng ml−1, MyBioSource, MBS143105) for 24 hours, as described previously [3].

In another approach, Amyloid beta 1-42 (Cayman Chemicals) was incubated at 37°C for 1 hour to form protofibrils and then added to astrocytes at a concentration of 1 µM on DIV13 for 24 hours.

### Antisense LNA Gapmers

Antisense LNA Gapmers targeting *Prdm16os* and *PRDM16-DT* and negative controls (NC) were designed and purchased from Qiagen having the following sequences: NC: GCTCCCTTCAATCCAA *Prdm16os*: TGCGACGTCTAAGATG *PRDM16-DT*: TACAGAACTGGTCATT

Transfection was performed using Lipofectamine RNAiMAX (ThermoFisher) according to the manufacturer’s instructions. Cells were treated on DIV12 and all experiments were performed on DIV14.

### *Prdm16os* overexpression using CRISPRa

Plasmids containing the guide RNA (gRNA), a dCas9-VP64 and a tGFP reporter gene were obtained from Origene (GE100074). The gRNAs for targeting Prdm16os were also designed by Origene and had the following sequences: gRNA-1: AGACGGTCACCTCGCCTCCA gRNA-2: GATAGTTGGGACACGGGTCC gRNA-3: GAGCCCGAAGCTGCAGCCAC As a negative control, a scramble gRNA (Scramble) with no target in the genome was used (GE100077). To increase gene expression, an enhancer vector (GE100056) was used in all cases.

The cells were transfected before seeding on DIV7 using the Neon Nxt electroporation device (ThermoFisher). 100,000 cells were electroporated using 0.5 µg CRISPRa plasmid and 0.1625 µg enhancer plasmid (1300 V, 20 ms, 2 pulses). Cells were seeded in 24 well plates and cultured for two days before treatment with the cytokine cocktail or PBS. RNA isolation and functional experiments were performed 24 hours later.

### Glutamate uptake

On DIV14, primary astrocytes were washed with HBSS for 10 min at 37°C. Then, they were incubated with 100 µM glutamate in HBSS for 1 hour at 37°C. Afterwards, the supernatant was collected and the amounts of remaining glutamate in the HBSS were measured using the Glutamate-Glo™ Assay (Promega) according to the manufacturer’s instructions. Luminescence was recorded with a FLUOstar® Omega (BMG).

### Lactate secretion

To estimate lactate secretion, the complete medium was changed one hour before performing transfection. After 48 hours, the medium was collected and diluted 1:40 in PBS. Lactate concentrations were measured using the Lactate-Glo™ Assay (Promega) according to the manufacturer’s protocol.

### Cytoplasmic and nuclear fractionation

Primary astrocytes cultured in 6 well plates were dissociated on DIV14 with 0.25% trypsin-EDTA. After centrifuging at 400 g for five minutes and washing the cells with PBS, 500 µL EZ prep lysis buffer (Sigma-Aldrich) supplemented with RNase inhibitor (Promega) was added to each sample and incubated on ice for seven minutes. The samples were centrifuged and the supernatant was aspirated and collected, containing the cytoplasmic fraction. The nuclear pellet was washed with 2 mL EZ prep lysis buffer, incubated on ice for five minutes and centrifuged again. The supernatant was aspirated and the pellet resuspended in 1.5 mL phosphate buffered saline (PBS) supplemented with 0.5% bovine serum albumin (BSA), RNase inhibitor and protease inhibitor (Roche). The samples were centrifuged and the supernatant aspirated, leaving 250 µL of liquid in the tube. TRIzol™ LS Reagent (ThermoFisher) was added to both the cytoplasmic and nuclear fraction and the samples were stored at -20°C until RNA was extracted.

### Glutamate treatment of primary neurons

Neuronal and neuron-astrocyte co-cultures were incubated on DIV 17 with 100 µM glutamate for 15 minutes at 37°C. Then, the medium containing glutamate was removed and replaced with fresh NB+ for 3 hours. Cell viability was measured using PrestoBlue (ThermoFisher).

### Spine density analysis

Dendritic spines were labelled using the dye 1,1 -dioctadecyl-3,3,3 ,3 - tetramethylindocarbocyanine perchlorate (Dil) (Life Technologies) as described previously [76], using 2% paraformaldehyde (PFA) for fixation. Spine density and dendrite length were measured with ImageJ software.

### MACS-sorting of oligodendrocytes, astrocytes and microglia

Cells were isolated from the cortex of three months old male C57B/6J mice using the adult brain dissociation kit (cat. no. 130-107-677, Miltenyi) according to the manufacturer’s protocol with minor modifications. Briefly, mice were sacrificed using pentobarbital and the brains were quickly removed. To remove major parts of the meninges, the brains were rolled over Whatman paper and then the cortices were dissected and placed into the enzyme mixes. The tissue was incubated at 37°C for 30 minutes in a water bath and triturated gently three times during this period. Then, the samples were applied to 40 µm cell strainers and the protocol was followed for debris and red blood cell removal. Oligodendrocytes were isolated using Anti-O4 microbeads (1:40, cat. no. 130-094-543), astrocytes using Anti-ACSA2 microbeads (1:10, cat. no. 130-097-678) and microglia with Anti-Cd11b microbeads (1:10, cat. no. 130-093-634). Purity of the cell type populations was determined by qPCR.

### In situ hybridization combined with immunofluorescence

We performed RNAscope Fluorescent Multiplex assays (ACD Bio) combined with immunofluorescence according to the manufacturer’s instructions for fresh frozen tissue sections. Briefly, 18 µm sections were fixed with 10% neutral buffered formalin (NBF) and dehydrated with ethanol. Hydrogen peroxide was applied to the sections, followed by the incubation with anti-Gfap (rabbit, Abcam; 1:250) at 4°C overnight. On the next day, after a post-primary fixation step with 10% NBF, the protocol for the RNAscope® Multiplex Fluorescent Reagent Kit v2 (Acd Bio) was followed, using probes designed against *Prdm16os* (Acdbio) and TSA Plus Cyanine 5 (Akoya Biosciences; 1:750) for detection of the probes. Afterwards, the slides were incubated with Alexa Fluor™ 555 goat anti-rabbit secondary antibody (1:1000, ThermoFisher) and DAPI (Sigma-Aldrich) and mounted using Prolong Gold Antifade Reagent (ThermoFisher). Images were acquired within a week after staining.

### Imaging

All images were taken with a Leica dmi8 microscope fitted with a STEDycon STED/Confocal (Abberior) in the confocal mode, with a 63X or 100X oil immersion objective.

### RNA extraction

Cells were lysed using TRI Reagent (Sigma-Aldrich) and total RNA was isolated using the RNA clean & concentrator-5 kit (Zymo Research) according to the manufacturer’s instructions.

RNA concentrations were determined by Nanodrop (ThermoFisher) or Qubit using the RNA HS Assay kit (ThermoFisher). The quality of the samples used for sequencing was assessed with a Bioanalyzer (Agilent Technologies).

### Library preparation and total RNA sequencing

Libraries were prepared using the SMARTer Stranded Total RNA Sample Prep Kit - HI Mammalian kit (Takara Bio) using 300 ng RNA as starting material. Libraries were amplified for 13 cycles and the quality of the preparations determined on the Bioanalyzer. The multiplexed libraries were sequenced in a NextSeq 2000 (Illumina) with a 50 bp single-read configuration.

### Bioinformatic analysis

Processing and demultiplexing of raw reads was performed using bcl2fastq (v2.20.2). For quality control of raw sequencing data, FastQC (v0.11.5) was used. Reads were aligned to the mouse (mm10) genome with the STAR aligner (v2.5.2b), and read counts were generated using feature Counts (v1.5.1). Differential gene expression was performed with DESeq2 (v1.38.3) [77] using normalized read counts and correcting for unwanted variation detected by RUVSeq (v1.32.0) [78]. GO term analysis was performed with clusterProfiler (v4.6.0) [30]. Analysis for the enrichment of transcription factors targeting downregulated genes was performed using ENRICHR (https://maayanlab.cloud/Enrichr/).

### cDNA, qPCR

cDNA was prepared with the Transcriptor cDNA first strand Synthesis Kit (Roche) using 20-800 ng RNA as starting material and random hexamer primers.

Synthesized cDNA was diluted up to ten-fold with nuclease-free water. qPCR reactions were prepared with Light Cycler 480 SYBR Master Mix (Roche) and run in duplicates in a Light Cycler 480 (Roche). Primer sequences can be found in supplemental table 3. Analysis was done using the 2^-DDCt method [79].

### Western Blot

Cells were lysed in RIPA buffer (ThermoFisher) containing protease inhibitor. Protein concentration was determined using the Pierce BCA Protein Assay Kit (ThermoFisher), and 20 µg were used per well of a 4–20% Mini-PROTEAN® TGX™ Precast Protein Gel (Bio-Rad). Protein denaturation was done using 8 M urea in Laemmli Sample Buffer (Bio-Rad) at 40°C for 60 minutes.. The gels were run at 90 V for 15 minutes, followed by 50 minutes at 120 V. For the transfer, low fluorescence PVDF membranes and the Trans-Blot Turbo Transfer System (both from Bio-Rad) were used.

Membranes were blocked with 5% BSA in PBS + 0.1% Tween-20 (PBS-T) for 1 hour at 4°C . Primary antibodies were incubated ON at 4°C in 1% milk in TBS-T or 5% BSA in PBS-T. The following primary antibodies were used: anti-Eaat1 (rabbit, Abcam; 1:2500), anti-Eaat2 (rabbit, Abcam; 1:800) and anti-Gapdh (mouse, ThermoFisher; 1:4000). On the next day, the membranes were washed with PBS-T and incubated with the respective fluorescent secondary antibodies (IRDye, LI-COR). After washing the membranes again, blots were imaged with an Odyssey DLx (LI-COR) and the images were analyzed with ImageJ software. Information on antibodies can be found in supplemental table 4.

### RNA immunoprecipitation

For RNA immunoprecipitation (RNA-IP), cells were cultured in T75 flasks. On DIV14, cells were dissociated with 0.25% trypsin-EDTA, washed with PBS and resuspended in fractionation buffer (10 mM Tris-HCl, 10 mM NaCl, 3 mM MgCl_2_, 0.5% Nonidet P-40 (NP-40), 1mM DTT, 100 units/mL RNase inhibitor, 1x protease inhibitor). The samples were incubated on ice for 10 minutes and then resuspended again to lyse the cells. After centrifuging for five minutes at 1000 g and 4°C, the supernatant, containing the cytoplasmic fraction, was transferred into fresh tubes. The nuclear fraction was rinsed with TSE buffer (10 mM Tris, 300 mM sucrose, 1 mM EDTA, 0.1% NP-40, 1mM DTT, 100 units/mL RNase inhibitor, 1x protease inhibitor) and centrifuged at 1000 g for five minutes. Then, the pellet was resuspended in fresh TSE buffer, transferred to bioruptor tubes and sonicated in a Bioruptor Plus (both Diagenode) for ten cycles (30s on, 30s off). The samples were incubated on ice for 20 minutes with occasional vortexing and centrifuged at 14500 g for ten minutes. The supernatant was transferred to a fresh tube and both fractions were flash-frozen and stored at - 80°C.

The nuclear lysates were thawed and the protein concentration determined using a BCA assay. 500 µg lysate per sample were pre-cleared using 25 µL Pierce™ Protein A/G magnetic beads (ThermoFisher) for 1 hour at 4°C to reduce unspecific binding. 1.5 µg per sample of anti-Rest (rabbit, ThermoFisher) antibody or the IgG isotype control (Abcam) was incubated with 50 µL Protein A/G magnetic beads for two hours at RT and washed with RNA-IP buffer (50 mM Tris-HCl, 100 mM NaCl, 32 mM NaF, 0.5% NP-40). Then, the samples, after taking 10% as input, were added to the beads and incubated overnight at 4°C with mixing. On the next day, the beads were washed five times with RNA-IP buffer, resuspended in proteinase K in proteinase K buffer (Qiagen) and incubated for one hour at 37°C. The supernatant was collected and RNA extracted using the RNA clean & concentrator- 5 kit (Zymo Research) according to the manufacturer’s instructions.

### Chromatin immunoprecipitation (ChIP)

Primary astrocytes were grown in 15 cm dishes and treated with NC or KD ASOs as described above. On DIV14, cells were dissociated with 0.25% trypsin-EDTA, washed with PBS and cross-linked using 1% PFA for 10 minutes. The reaction was quenched with 125 mM glycine for 5 minutes. The pellets were washed with PBS and, after removing all the liquid, flash-frozen in liquid-nitrogen and stored at -80°C. For chromatin shearing, the weight of each pellet was measured and the pellets were resuspended in a 10x volume of RIPA buffer supplemented with 0.8% SDS and 1x protease inhibitor. The samples were sonicated for 20 cycles in a Bioruptor plus (30s on, 45s off). Chromatin shearing was checked by taking a small aliquot and decrosslinking the DNA by RNase A and Proteinase K treatment for 1 hour at 65°C. DNA was isolated using the ZYMO ChIP Clean and Concentrator Kit. Sheared chromatin size was determined using Bioanalyzer 2100 (DNA high sensitivity kit) and the concentration was measured using Qubit 2.0 fluorometer (DNA high sensitivity kit).10 µg of chromatin was used along with 3 µg of antibody to do ChIP for Rest (rabbit, ThermoFisher) or the IgG isotype control (Abcam). For the H3K27me3 mark, 800 ng of chromatin and 2 µg of H3K27me3 antibody (rabbit, MerckMillipore) was used. ChIP was performed as described previously [80] with minor modifications. After pre-clearing the samples and overnight incubation with the respective antibodies, 25 μL of BSA-blocked protein A magnetic beads were added to each sample and the mixture was incubated on a rotator at 4 °C for 2 h. The complexes were washed with Low Salt Wash buffer (20 mM Tris-HCl at pH 8, 150 mM NaCl, 1% Triton X-100, 2 mM EDTA, 0.1% SDS, Roche Complete protease inhibitors), High Salt Wash buffer (20 mM Tris-HCl at pH 8, 500 mM NaCl, 1% Triton X-100, 2 mM EDTA, 0.1% SDS, Roche Complete protease inhibitors), lithium chloride (LiCl) Wash buffer (10 mM Tris-HCl at pH 8, 1% NP-40, 1% sodium deoxycholate, 1 mM EDTA, 250 mM LiCl) and 1x Tris-EDTA (TE) buffer. The DNA was eluted from the beads and purified using the ZYMO ChIP Clean and Concentrator Kit. The DNA was eluted with 50 µL elution buffer and 1 µL was used for the qPCR.

### Statistical analysis

Statistical analysis was done using GraphPad Prism version 9. All graphs are shown as mean ± SEM unless stated otherwise. For data analysis, either a two-tailed unpaired t-test or a one-way ANOVA with Tukey’s post hoc test were applied. Enriched gene ontology and pathway analysis was performed using Fisher’s exact test followed by a Benjamini-Hochberg correction.

## Acknowledgements

AF was supported by the DFG (Deutsche Forschungsgemeinschaft) priority program 1738, SFB1286 and GRK2824;The German Federal Ministry of Science and Education (BMBF) via the ERA-NET Neuron project EPINEURODEVO; The EU Joint Programme- Neurodegenerative Diseases (JPND) – EPI-3E; Germany’s Excellence Strategy - EXC 2067/1 390729940. FS was supported by the GoBIO project miRassay (16LW0055) by the German Federal Ministry of Science and Education (F Sananbenesi, A Schutz). AF and ID are supported by NIH RF1AG078299.

## Data availability

RNA-sequencing data will be available via GEO and EGA.

---

**Fig S1.**
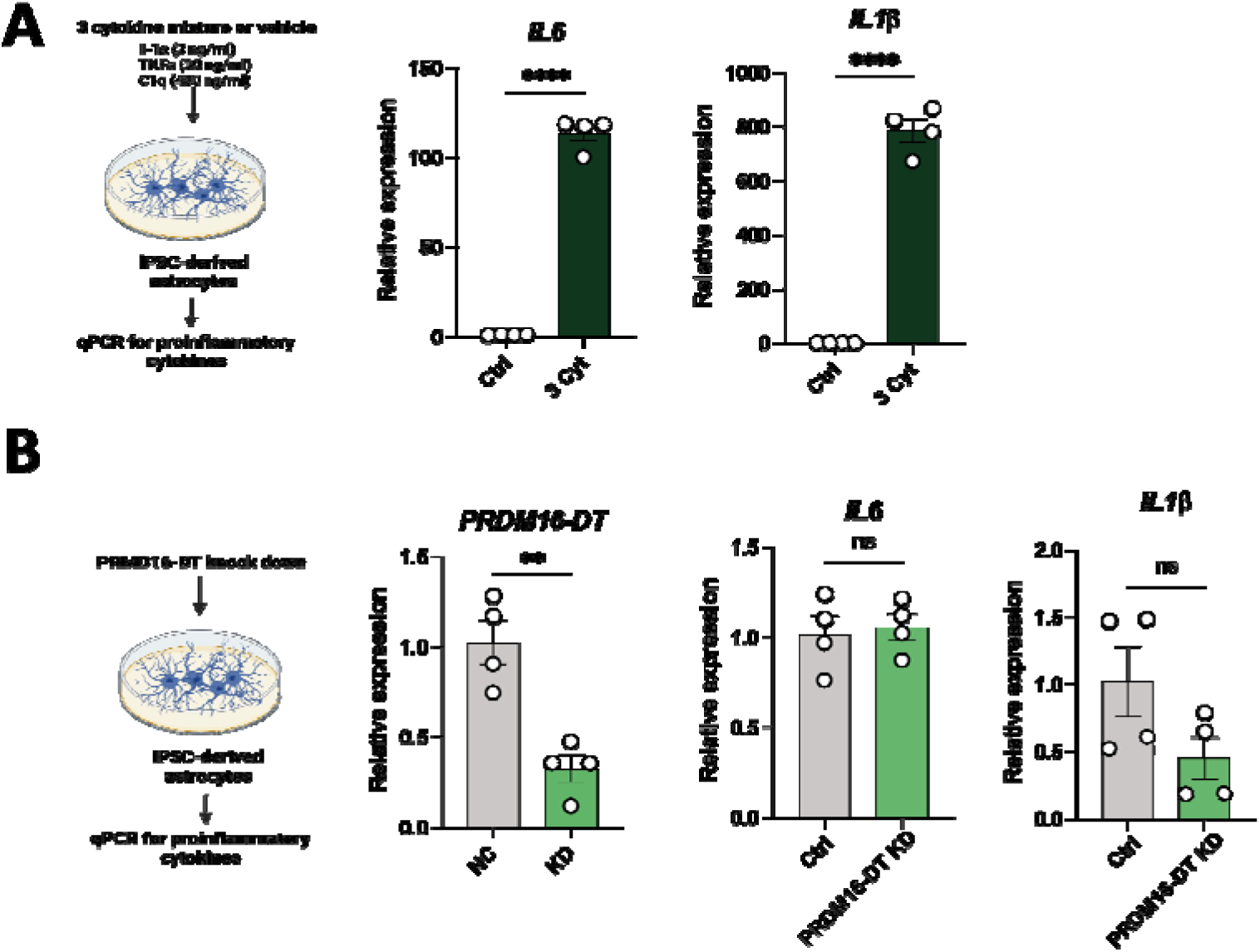

